# The best of two worlds: Decoding and source-reconstructing M/EEG oscillatory activity with a unique model

**DOI:** 10.1101/2023.03.24.534080

**Authors:** Britta U. Westner, Jean-Rémi King

## Abstract

The application of decoding models to electrophysiological data has become standard practice in neuroscience. The use of such methods on sensor space data can, however, limit the interpretability of the results, since brain sources cannot be readily estimated from the decoding of sensor space responses. Here, we propose a new method that combines the common spatial patterns (CSP) algorithm with beamformer source reconstruction for the decoding of oscillatory activity. We compare this method to sensor and source space decoding and show that it performs equally well as source space decoding with respect to both decoding accuracy and source localization without the extensive computational cost. We confirm our simulation results on a real MEG data set. In conclusion, our proposed method performs as good as source space decoding, is highly interpretable in the spatial domain, and has low computational cost.

## 1 Introduction

In the last two decades, decoding has become a go-to method to predict neural representations from high-dimensional recordings of brain activity (Cichy et al., 2017; King et al., 2018; Kriegeskorte, 2015; Pantazis, 2020; Pantazis et al., 2018; van Vliet et al., 2017). However, decoding often faces a major limitation: interpretability. In particular, *spatial decoders* are trained across multiple sensors (electrophysiological data) or multiple voxels (magnetic resonance data) to identify *whether* various neural signals differ from one another, but not *how* this difference is characterized in terms of brain activity (de Cheveigné and Simon, 2008). When applied to whole-brain recordings, this issue can become highly problematic. In magneto- or electroencephalography (M/EEG), linear decoders are often trained on the sensor-level data (e.g., Blom et al., 2020; Foster et al., 2021; King et al., 2016; Konvalinka et al., 2014; Olivetti et al., 2014), and it can thus be challenging to know the source of the decoded information (King et al., 2018; Kriegeskorte and Douglas, 2019).

One reason for this is the fact that the coefficients of linear models are not directly interpretable because they do not reflect informative brain activity. Haufe and colleagues introduced an analytical method to obtain the unique and interpretable brain activity “pattern” that is associated with the coefficients of the linear model. However, while this method increases interpretability as compared to the model coefficients, there is still a drawback to this method. The resulting patterns are still in sensor space if the decoding model was computed on sensor space data. This limits the interpretability of the underlying sources through sensor or electrode positions, especially across participants. Moreover, the nature of the patterns does not allow for post-hoc source reconstruction: the positive and negative values now refer to classes and not electromagnetic sources and sinks. How this changes the structure of the data is illustrated in Fig. 1. One possible solution to the interpretability problem is the application of decoding models in source space. This increases interpretability and might also increase decoding performance since source reconstruction methods unmix the sensor signals and thus can act as a denoising step (Westner et al., 2022). Furthermore, it can enable decoding across participants since the data can be represented in a common source space (Westner et al., 2018). However, this adds considerable computational cost due to the increase in data points (thousands of source points as compare to hundreds of sensors).

**Figure 1:**
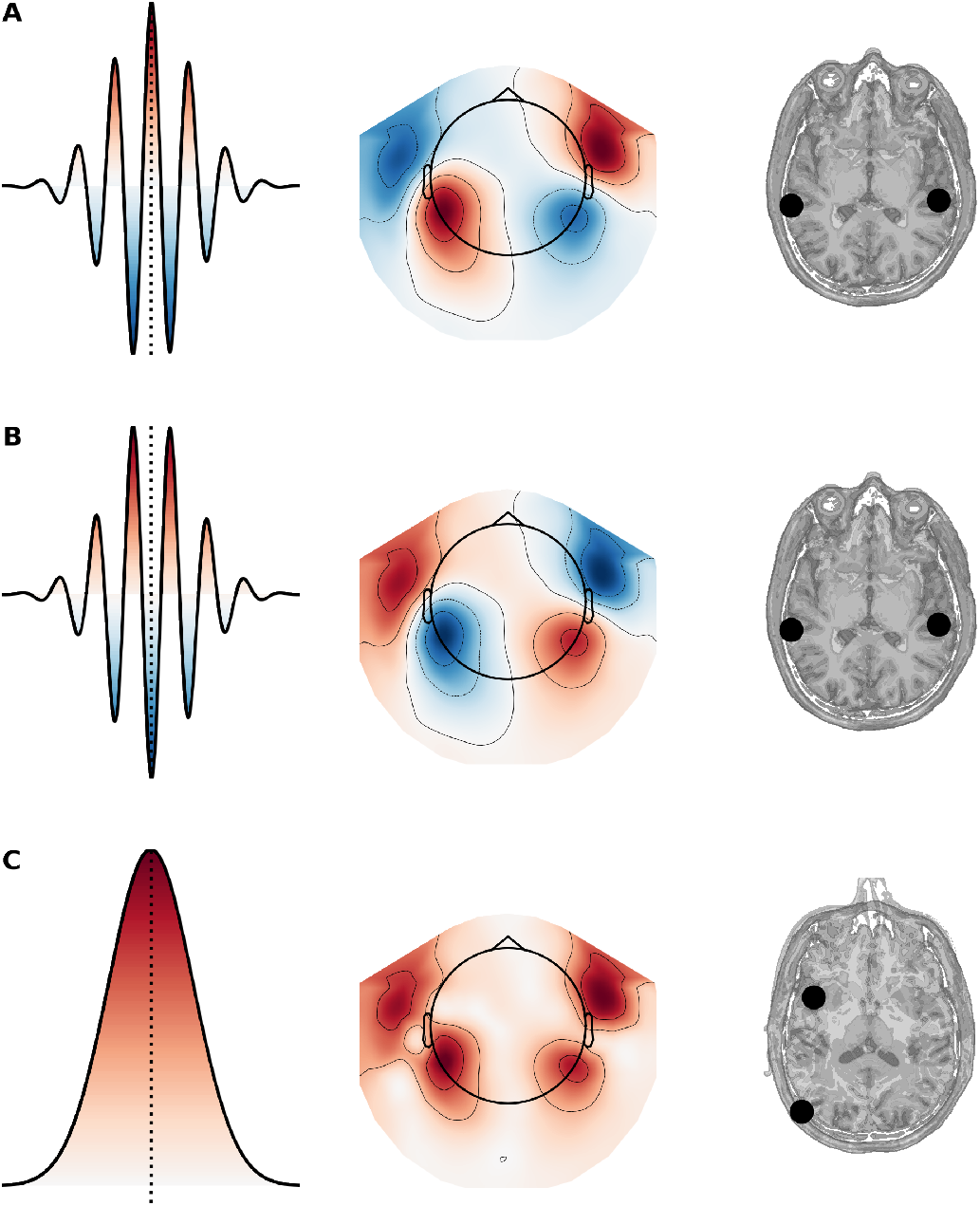
Incompatibility of sensor space decoding patterns and source reconstruction A and B. Oscillatory activtiy, left to right: oscillation, topography, and source reconstruction. **C** Decoding pattern of the activity in A or B: The sources are misplaced, since the sensor pattern does not reflect sources and sinks anymore, but the importance of the underlying brain activity for the decoding.

In this paper, we compare the performance of sensor and source space decoding, but also present a new approach for the decoding of oscillatory activity. We propose to decode induced activity using the common spatial pattern algorithm (CSP; Koles, 1991; Koles and Soong, 1998; Ramoser et al., 2000) in combination with beamformer source reconstruction. CSP is a linear decoding method which aims at finding a set of orthogonal components such that the variance onto which M/EEG signals are projected maximally discriminates two conditions (see Eq. (6) and Eq. (7) in Methods).

We can formalize three distinct pipelines which differ especially with respect to the order of operations in the pipeline:

1. **power-first pipeline**: Power estimation → Linear classifier → Haufe’s patterns → Source reconstruction
2. **source-first pipeline**: Source modeling → Power estimation → Linear classifier → Haufe’s patterns
3. **CSP-first pipeline**: CSP → Linear classifier → Haufe’s patterns → Source reconstruction

We predict that the power-first pipeline will fail in terms of source localization because the initial power estimation introduces an irreversible and nonlinear transformation of M/EEG activity. We predict that the source-first pipeline will work well, but will also present a large computational cost, because there are many more brain source points than MEG sensors. Finally, we predict that our newly proposed CSP-first pipeline will be both valid and computationally efficient.

To compare these alternatives, we synthesize brain signals and evaluate the three pipelines on their (1) sensitivity (i.e., how good the decoding performance is), (2) interpretability (how precise their source localization is) and (3) computational load. We also evaluate our method on a public MEG data set. Our analyses show that our CSP-first pipeline successfully and efficiently combines CSP and a linear classifier while ensuring a valid source modeling of its coefficients.

## 2 Methods

### 2.1 Simulations

We simulate oscillatory brain activity data following the experimental design of a real public MEG data set (the so-called `sample` data set in MNE-Python, Gramfort et al. (2013)). Contrary to real MEG data, synthetic data possess a ground truth for source localization, and thus is well-suited for the evaluation of different inverse modeling pipelines.

Specifically, we synthesize two distinct data sets which vary in terms of difficulty of decoding and source localization. Specifically, we synthesize (1) a visual and an auditory source in the corresponding distant cortices and (2) two distinct visual sources within the left and right occipital cortices, respectively. The objective of the decoders for each data set is to classify single trials of oscillatory activity into the two classes: (1) *visual-audio* or (2) visual left and visual right, *left-right*.

The MEG data are synthesized by placing a source in a randomly chosen voxel within a predefined region (see below) of a surface-based grid. The corresponding neural activity of this source is simulated with a sinusoid oscillating at a frequency of either 72 Hz or 75 Hz, depending on the class. To limit edge artefacts which would leak to multiple frequencies, the amplitude of these oscillations is attenuated at the edges of each trial using a Gaussian taper, resulting in an active window from 450 to 550 ms, centered in the 1 s long trial. One hundred trials are generated for each class. To make sure the decoders cannot make use of evoked responses, the phase of the simulated oscillatory activity is randomized across trials. The data is then projected to sensor space with a standard surface forward model. The sensor-space signal-to-noise ratio (SNR) then is manipulated by adding white noise (between −45 and 5 dB in 5 dB steps). The noise level is determined on the first condition and then applied to the second one, such that the conditions are not distinguishable based on a difference in the noise alone, e.g., during the baseline period. Thus, the sensor data **B** was generated as follows:

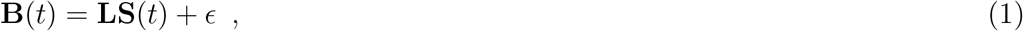

where **L** ∈ ℝ^*M×N*^ is the forward model for *M* sensors and *N* source points, **S**(*t*) ∈ ℝ^*N×*1^ is the simulated source-level activity at time point *t*, and *ϵ* ∈ ℝ^*M×*1^ is the vector of random white noise added to the sensor-level data. The source-level activity of an active source **S**_*active*_(*r, t*) at source point *r* and time point *t* was simulated as

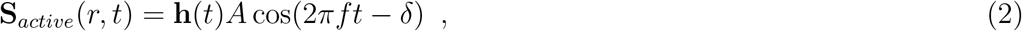

with **h** ∈ ℝ^1*×T*^ being the vector of a Hanning taper for a time series of length *T, A* referring to the amplitude, *f* being the frequency, and *δ* the phase of the oscillation.

For both contrasts, *visual-audio* and *left-right*, 200 simulations are generated with variable source i) location and ii) orientation of the two sources. The position of the sources is selected within a region of interest defined by Freesurfer’s aparc parcellation (Fischl, 2004), for the *visual-audio* contrast to “visual left hemisphere” and “auditory right hemisphere”, for the *left-right* contrast to “visual left hemisphere” and “visual right hemisphere”.

We report the means and standard deviations of decoding performance, localization error, and source spread across the 200 runs and for each input SNR.

### 2.2 Decoding

Decoding is based on an *ℓ*_2_-regularized logistic regression:

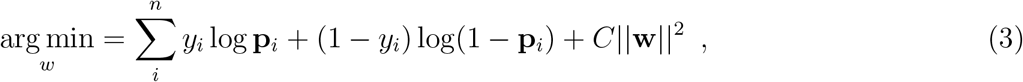

where 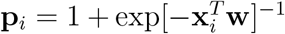 constitutes a vector of *c* MEG channels at trial *i*, **w** ∈ ℝ^*c*^ is the decoder’s coefficients, *y*_*i*_ ∈ [−1, 1] codes for the category of the trial (e.g., visual versus auditory), and *C* is the regularization coefficient. We train the logistic regression with scikit-learn (Pedregosa et al., 2011). Except if stated otherwise, all model hyperparameters are kept to their default MNE-Python values (i.e., intercept *liblinear* solver, *C* = 1.0).

#### Power-first pipeline

The single-trial power estimates are computed over 0–700 ms by first bandpass-filtering (75–95 Hz) and then squaring the data. The values are then standardized and scaled before being used as input to a logistic regression.

#### Source-first pipeline

The epoched data are first projected to source space using a unit-noise-gain linearly constrained minimum variance (LCMV) beamformer with a forward model based on a volumetric grid. The data covariance matrix is computed on the active 700 ms of the data and regularized with 1 % of the sensor power. Prewhitening is applied using a noise covariance matrix computed on a 200 ms baseline period (800–1000 ms). The source orientation is chosen to maximize spatial filter output power. The unit-noise gain scalar LCMV spatial filter weights **w** ∈ ℝ^*M×*1^ for a source point *r* and its optimal source orientation are computed as follows:

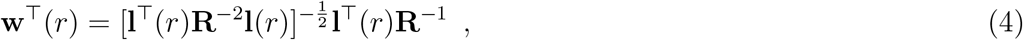

with **l**(*r*) ∈ ℝ^*M×*1^ being the forward model for this source point and **R** ∈ ℝ^*M×M*^ being the covariance matrix of the data. The filter weights **w** are then applied to the sensor space data **b** to estimate source activity ŝ at source point *r* and time point *t*:

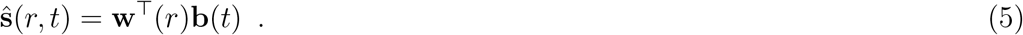

The power of the source space data is then computed per trial over the 700 ms active window. The data are standardized and scaled and fed to the logistic regression algorithm.

#### CSP-first pipeline

The epoched sensor data are standardadized and scaled across time for each sensor, projected through a 4-dimensional CSP operator, and finally used as input to a logistic regression.

The CSP procedure finds a spatial filter **W** that decomposes the data into orthogonal components (Koles, 1991). This is achieved through the simultaneous diagonalization of the two covariance matrices of the data with *M* channels, **R**^+^ ∈ ℝ^*M×M*^ and **R**^−^ ∈ ℝ^*M×M*^, which belong to the two classes (Blankertz et al., 2008; Koles, 1991). This simultaneous diagonalization can be achieved through a generalized eigenvalue decomposition, such that the following is satisfied:

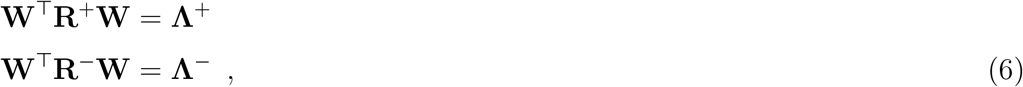

with **W** ∈ ℝ^*M×M*^ being the spatial filter and **Λ**^*c*^ ∈ ℝ^*M×M*^ being a diagonal matrix whose elements are obtained through the generalized eigenvalue decomposition

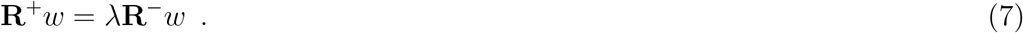

The generalized eigenvectors **w** give the columns of the spatial filter **W**, and the diagonal of Λ^*c*^ in Eq. (6) is given by *λ* = *λ*^+^*/λ*^−^ (cf. Eq. (7)).

#### Projection of sensor space patterns to source space

To make the output interpretable, decoding patterns are derived from the fitted models following Haufe et al. (2014). The patterns of the forward model **A** ∈ ℝ^*M×K*^ for *M* measurement channels and *K* latent factors are obtained by:

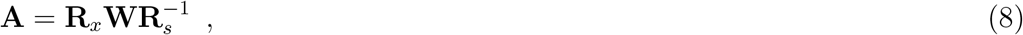

with **R**_*x*_ ∈ ℝ^*M×M*^ being the covariance matrix of the data, **R**_*s*_ ∈ ℝ^*K×K*^ being the covariance matrix of the latent factors, and **W** ∈ ℝ^*M×K*^ being the filters of the decoding model.

The obtained sensor space patterns are projected to source space, both for the power-first and the CSP-first pipeline. To this end, a beamformer spatial filter is constructed using the original, bandpass-filtered data for the data and noise covariance matrices. The constructed spatial filter has the same specifications as the beamformer for the source space decoding described above. The sensor space pattern is whitened using the whitener from the construction of the spatial filter. Then, the spatial filter is applied to the whitened pattern to create a source reconstruction of this pattern. In the case of the CSP pipeline, the four obtained patterns (one per component) are first combined by multiplying them with their respective logistic regression weights, then summed across all components, and then whitened and projected through the beamformer spatial filter. Since the optimal source orientation obtained with LCMV beamformers is arbitrary by 180°, the reconstructed pattern can look patchy with its arbitrary positive and negative source estimates, which can make interpretation hard especially if the two classes are close to each other in space. We therefore combined the patterns and components first and then beamformed and multiplied each of them with either -1 or 1 before summing over the patterns in source space.

### 2.3 Evaluation

We evaluate the methods on their decoding performance as well as on their interpretability (i.e., spatial precision). To this end, we compare the obtained source patterns across methods regarding (1) localization precision and (2) source spread. Finally, we apply our three decoding pipelines to a real MEG data set. Additionally, to investigate the role of the regularization parameter *C* in the logistic regression, we perform a grid-search for *C*. The value for *C* that optimizes the decoding performance is searched among 10 values between 0.0001 and 1000, evenly spaced on a logarithmic scale. This is done for all three decoding pipelines and 200 simulations, but only one of the two contrasts (the *visual-audio* contrast).

#### Decoding performances

All decoding approaches are embedded in a 5-fold stratified cross-validation which includes the whole decoding pipeline. For the source-first pipeline this means that the computation of the beamforming weights are also crossvalidated, i.e., the weights are computed with a covariance matrix that is estimated based on the training data set. To evaluate decoding performance, the area under the curve of the receiver operating characteristic (ROC AUC) is computed and the chance level is calculated for *p* = 0.05 using a binomial distribution (cf. Combrisson and Jerbi, 2015).

#### Spatial precision

The obtained “source patterns” are evaluated regarding their localization error and spread. Localization errors are computed as the Euclidean distance between the simulated source and the peak voxel location of the reconstructed pattern. The spread of the source *g* is computed as the weighted voxel count of all activity *s*(*x*) greater than 70 % of the maximum activity *s*_*max*_, with voxels being weighted based on their distance from the peak location:

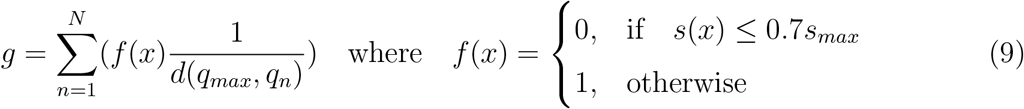

Here, *d*(*q*_*max*_, *q*_*n*_) denotes the Euclidian distance between the peak voxel *q*_*m*_*ax* and the current voxel *q*_*n*_. Higher values represent a wide spread of neural response around the peak, whereas small values represent a sharp peak response.

### 2.4 Real data

To validate our simulation results with real data, we test our analyses on the SPM faces data set as distributed with MNE-Python. This MEG face perception data set is openly available as the “Multi-modal Face Dataset” from the SPM software (https://www.fil.ion.ucl.ac.uk/spm/data/mmfaces/). During the recording, faces and scrambled faces were presented to the participant in random order (cf. Henson et al., 2003). We aimed to decode the high-frequency activity induced by faces versus scrambled faces, and based the choice for the processing parameters on the face-processing literature (Dobel et al., 2011; Itier and Taylor, 2004; Liu et al., 2018; Zion-Golumbic and Bentin, 2007). Specifically, the MEG data is filtered between 60–95 Hz to isolate oscillatory activity in the gamma band that is not dominated by the event-related fields and not influenced by line noise. The power of this oscillatory activity is then taken for a window from 180–400 ms post stimulus to further avoid any smeared activity from the M170 peak for face processing. Both the power-first decoding pipeline and the CSP-first pipeline are trained on the sensor space data. For the source-first pipeline, the data are first source-reconstructed using a unit-noise-gain LCMV beamformer with the same specifications as described for the simulations above. The corresponding data covariance matrix is computed on the window of 150–450 ms after stimulus presentation and regularized with 1 % of the sensor power, whereas the noise covariance matrix is constructed from the baseline (450–150 ms prior to stimulus presentation). The chance level for *p* = 0.05 is computed using a binomial distribution. All MEG analyses are done using the open-source toolbox MNE-Python (Gramfort et al., 2013). Analysis code can be found at https://github.com/britta-wstnr/source_decoding.

## 3 Results

### 3.1 Simulation results

#### Decoding scores

The results from the simulation show that the source-first pipeline outperforms the power-first pipeline for low input SNRs (Fig. 2A): while the source-first pipeline can distinguish classes significantly better than chance for an SNR of −30 dB for both *visual-audio* and *left-right*, the power-first pipeline only reaches successful classification above chance level at an input SNR of −25 dB. Likewise, a perfect separation of both classes with an ROC AUC of 1.0 is reached with lower input SNRs for the source-first pipeline (−20 dB for both *visual-audio* and *left-right*) than for the power-first pipeline (−15 dB for both contrasts). Critically, our CSP-first pipeline performs comparably to the source-first pipeline.

**Figure 2:**
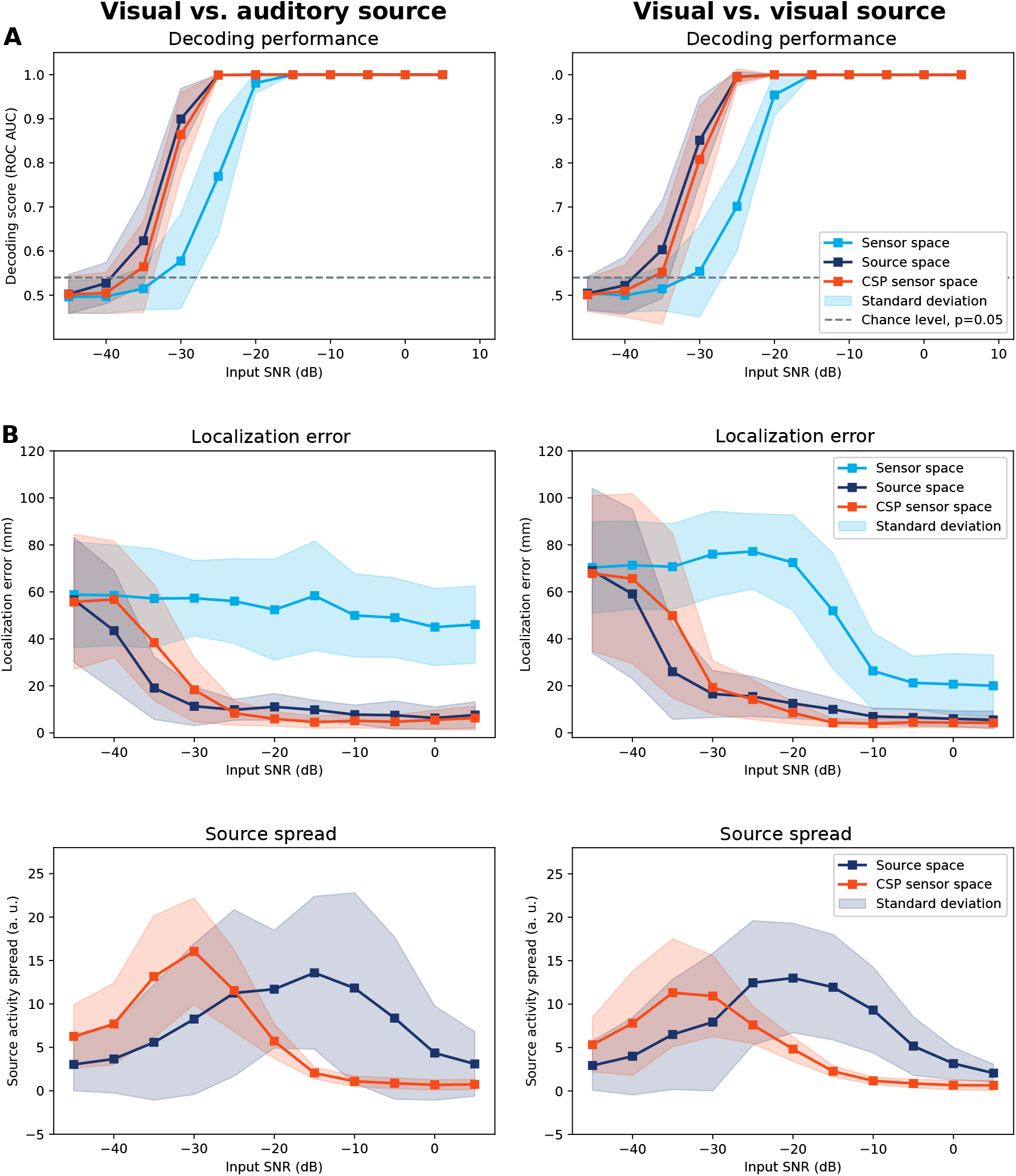
Simulation results. Results for visual vs. auditory sources (*visual-audio*, left column) and visual vs. visual sources (*left-right*, right column). Shaded areas depict standard deviations. **A** Decoding performance (ROC Area under curve) for different input SNRs, comparing the power-first pipeline (light blue), source-first pipeline (dark blue), and CSP-first pipeline (orange) decoding. **B** Localization error of the source peak with respect to the simulated source (ground truth) for power-first pipeline pattern (light blue), the source-first pipeline pattern (dark blue), and the beamformed CSP-first pipeline pattern (orange). **C** Source spread of the source-first pipeline (dark blue) and beamformed CSP-first pipeline pattern (orange) as a function of input SNR.

#### Localization error and source representation

How do these three pipelines compare in terms of source localization? To address this issue, we first estimated the decoding patterns of the power-first and CSP-first pipelines, and then beamformed them. For the source-first pipeline, we compute the decoding pattern in source space. Finally, we compare these three estimates to the ground truth (Fig. 2B).

The power-first pipeline clearly fails for the *visual-audio* case (Fig. 2B, left), while the localization error decreases with increasing input SNR for the *left-right* case (Fig. 2B, right). Even in the more favorable *left-right* case, the localization error still amounts to 20.00 mm (average; standard deviation: 13.09) for an input SNR of 5 dB (also compare Table 2 in the Supplementary Material).

By contrast, the localization errors of the source-first and the CSP-first pipelines decrease with increasing input SNR for both of the two classification problems. There is no clear difference in localization accuracy between these two approaches for either decoding problem (compare also Table 1 and 2 in the supplementary material).

**Table 1:** Localization errors visual-audio.

| Input SNR (dB) | -45 | -40 | -35 | -30 | -25 | -20 | -15 | -10 | -5 | 0 | 5 |
| --- | --- | --- | --- | --- | --- | --- | --- | --- | --- | --- | --- |
| **Sensor space decoding** |  |  |  |  |  |  |  |  |  |  |  |
| mean | 58.83 | 58.52 | 57.20 | 57.32 | 56.06 | 52.41 | 58.34 | 50.00 | 49.03 | 45.01 | 46.11 |
| std | 22.50 | 21.37 | 21.14 | 16.01 | 18.03 | 21.44 | 23.28 | 17.76 | 16.94 | 16.39 | 16.40 |
| **Source space decoding** |  |  |  |  |  |  |  |  |  |  |  |
| mean | 56.59 | 43.49 | 19.04 | 11.35 | 9.83 | 11.09 | 9.78 | 7.76 | 7.52 | 6.33 | 7.60 |
| std | 26.54 | 25.54 | 13.35 | 8.26 | 4.45 | 5.64 | 4.09 | 4.05 | 6.02 | 4.73 | 5.59 |
| **Sensor space decoding with CSP** |  |  |  |  |  |  |  |  |  |  |  |
| mean | 55.74 | 56.79 | 38.40 | 18.27 | 8.38 | 5.95 | 4.56 | 5.14 | 4.74 | 5.50 | 6.32 |
| std | 28.79 | 24.71 | 24.75 | 13.56 | 5.12 | 3.08 | 2.54 | 3.05 | 2.87 | 4.14 | 5.07 |

**Table 2:**
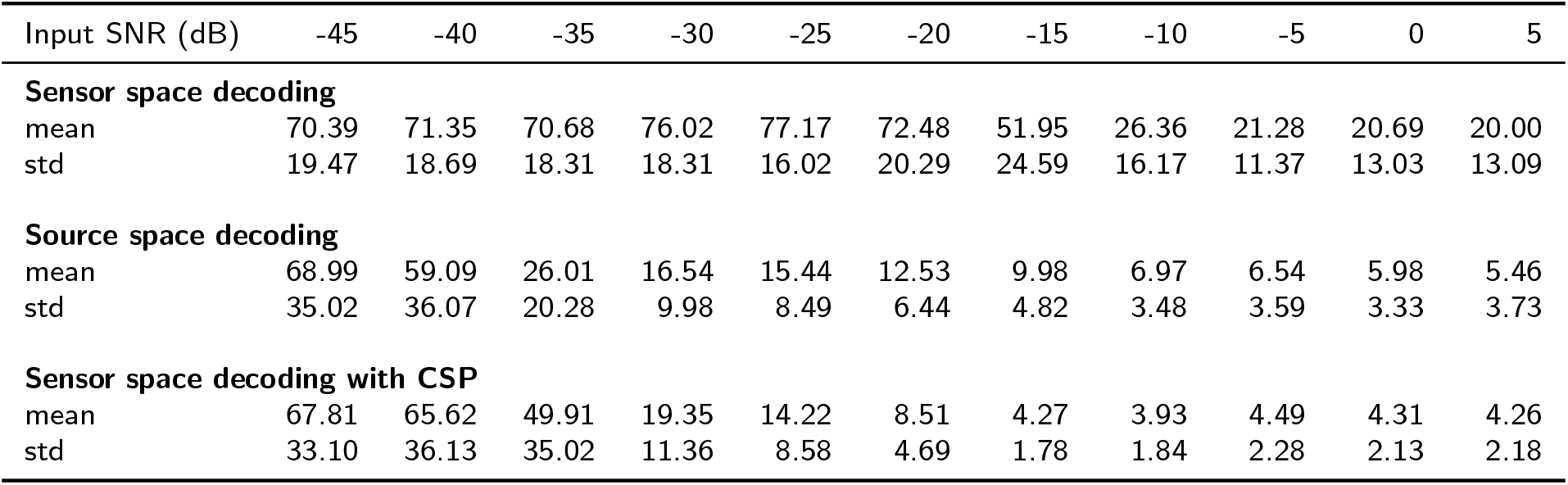
Localization errors left-right.

Fig. 3 shows exemplary source space patterns for an input SNR of −10 dB and one of the 200 realizations. As Fig. 2B suggests, both the source-first and the CSP-first pipelines reconstruct source patterns with a close match to the ground truth, while the power-first pipeline fails. The supplementary material holds a similar visualization for the *visual-audio* case (Fig. S1).

**Figure 3:**
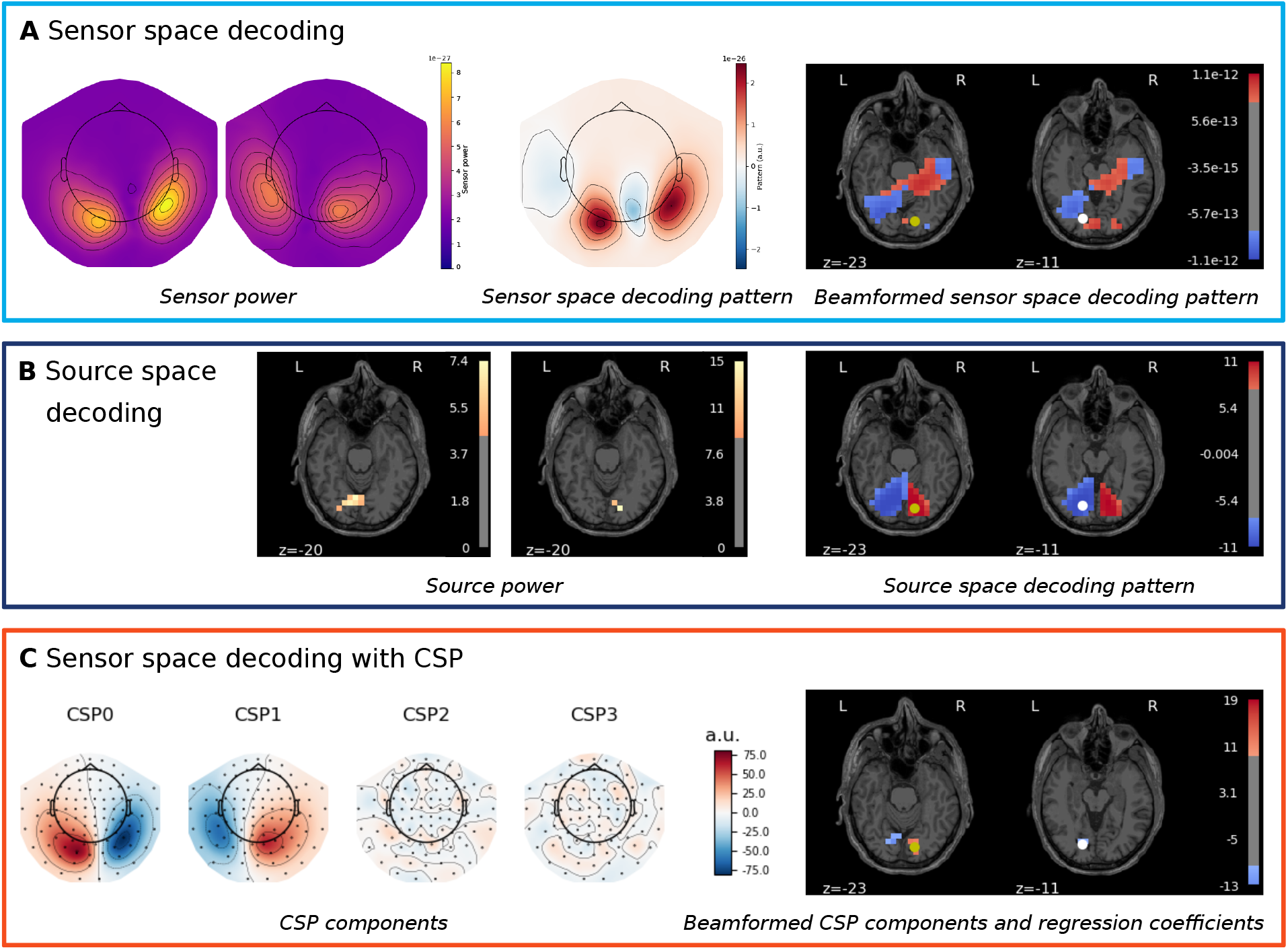
Visualization of decoding patterns for the simulation. The example corresponds to the *left-right* contrast and an input SNR of −10 dB. In all source plots, the true sources for the two classes are marked with a yellow and white dot. **A** Power-first pipeline. Sensor space power (left), sensor space decoding pattern after logistic regression (middle) and source space projection (right). **B** Source-first pipeline. Source space power (left) and source space decoding pattern (right). **C** CSP-first pipeline. CSP components (left) and the beamformed source estimate of the combined CSP components and logistic regression coefficients (right).

For the source-first and CSP-first pipelines, we also evaluated the spread of the source pattern. Figure 2C shows that the source estimates of the CSP-first pipeline are considerably sparser and less variable than those of the source-first pipeline at high input SNRs. This can be explained by the beamforming of the orthogonal components in the CSP-first pipeline - as opposed to the noisier raw power data in the source-first case.

#### Optimal regularization

The results reported above were obtained with a fixed regularization parameter of *C* = 1.0 for the logistic regression. To ensure that this arbitrary choice did not skew the results in favor of a particular decoding approach, we performed a grid search for the regularization parameter. Figure 4A shows the decoding performance for the optimal regularization parameter compared to the fixed parameter of *C* = 1.0 over 200 realizations. The chosen regularization parameter *C* = 1.0 performs comparably to the optimal regularization over 200 runs for all three approaches. As expected, Figure 4B shows that the optimal parameter exhibits a high variance for lower input SNRs, while a regularization with the lowest parameter of *C* = 0.0001 seems to be optimal for higher input SNRs.

**Figure 4:**
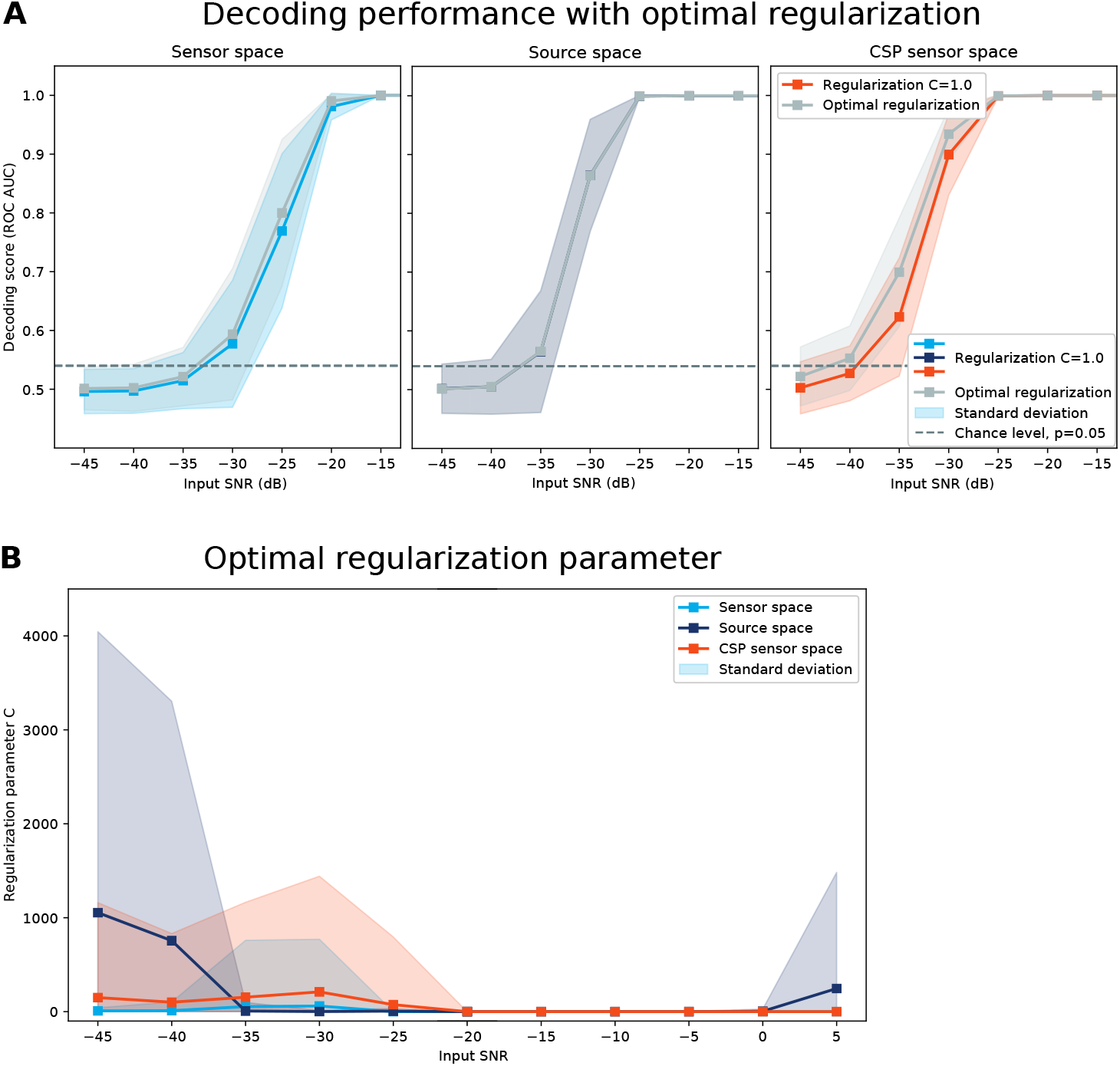
Grid search results. Search for the optimal regularization parameter *C* for the logistic regression. **A** Decoding scores for optimal regularization for each of the 200 realizations (light blue for sensor space, dark blue for source space, and orange for CSP sensor space decoding) compared to *C* = 1.0 for all 200 realizations (grey). **B** Optimal regularization values averaged across simulations. All simulations for the optimization of *C* are done for the auditory-visual contrast.

#### Computational time

Analyses were done on a high-performance computing cluster, and simulation realizations were distributed across different computing nodes. Unsurprisingly, the power-first pipeline proved to have the lowest computational time with an average time of 0.07 s across the 200 simulation runs (Fig. 5). This result is expected because this pipeline computes both the decoding models and the patterns based on sensor-space data which is fairly low-dimensional. By contrast, the source-first pipeline appears to be the slowest approach with an average time of 156.95 s. This pipeline not only uses a lot more features for the decoding (source points instead of sensors), but also has to estimate the beamformer spatial filters. Finally, our CSP-first pipeline took on average 14.06 s and is thus more than 10 times faster than the source space decoding (not including the projection of the pattern to source space). Here, decoding is done in sensor-space, but the pipeline also incorporates the estimation of the CSP components, which takes up additional time.

**Figure 5:**
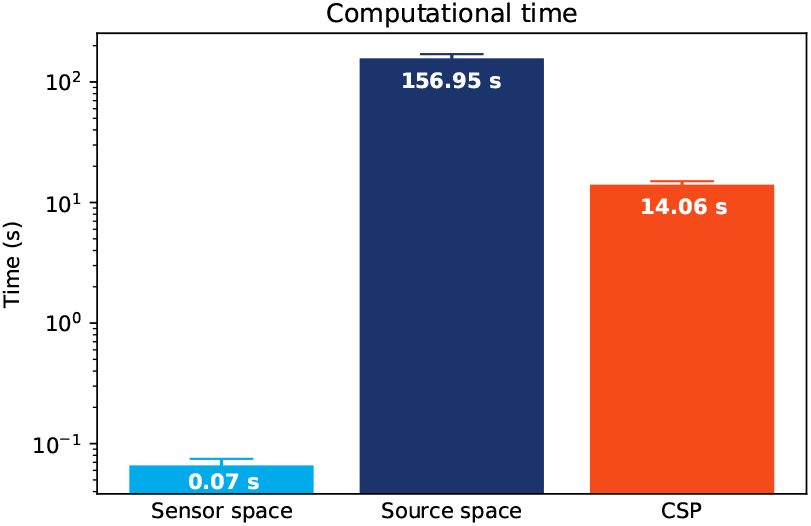
Computation time. Computational time averaged across the 200 simulations for visual versus auditory source. Note the logarithmic y-axis.

### 3.2 Real data results

We validated our simulation results on a real data set, classifying the percept of faces and scrambled faces using high frequency oscillatory activity. This is a relatively difficult classification problem, because the SNR of single trial gamma activity is notoriously low (Westner et al., 2018). Fig. 6A shows that both the source space and CSP sensor space classification are significantly better than chance across the five cross-validation folds (*p <* 0.05, as determine by a binomial test, cf. Combrisson and Jerbi (2015)): the source space approach and the CSP sensor space approach reach an ROC AUC of 0.6012 and 0.5859, respectively. The sensor space logistic regression does not perform above chance level with an ROC AUC of 0.4779.

**Figure 6:**
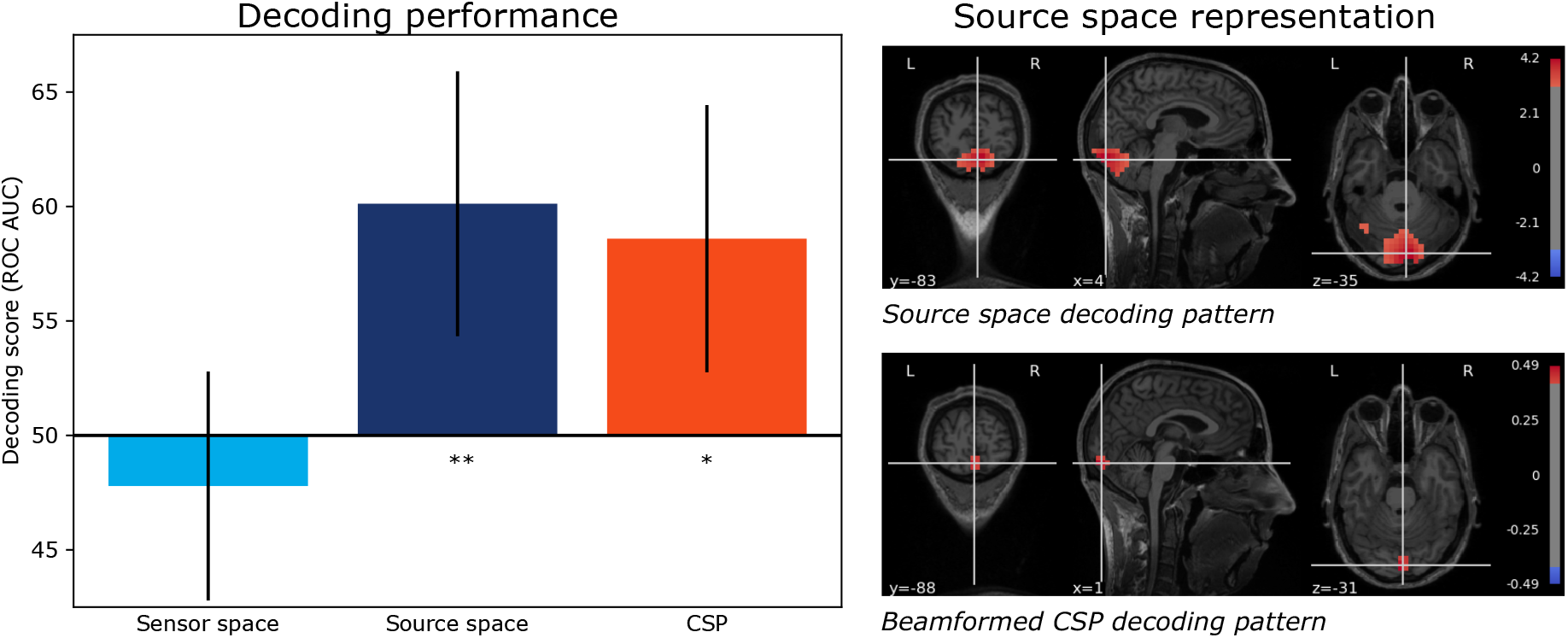
Real data results. Classification of faces vs. scrambled faces using high frequency activity. **A** Decoding performance (ROC AUC). Error-bars denote standard deviation across cross-validation folds. Asteriks refer to significance as obtained with a binomial test against chance level. **B** Decoding pattern for source decoding (upper panel) and beamformed sensor decoding pattern for the CSP decoding approach (lower panel). Both patterns are shown at a threshold of 75 % of maximal absolute activity.

Figure 6B shows the source representations of the classifier patterns. The top brain plot shows the source space decoding pattern with a clear peak in visual cortex. The beamformed CSP pattern matches this result closely, the peak being localized only a few voxels from where the maximum falls in the source space decoding case. As with the simulations, the beamformed CSP pipeline results appears more focal than the source decoding pattern.

These results corroborate our analyses of the synthetic data: our CSP-first pipeline appears to be equivalent with the source-first pipeline regarding both the decoding performance and source reconstruction, and can yet be computed over ten times faster.

## 4 Discussion

In this study, we compare three analytical pipelines designed to both decode and source-localize MEG oscillatory activity.

Overall, the source-first pipeline outperforms the power-first pipeline both in terms of classification accuracy and source localization. These results are expected: Source-localization is invalid in the power-first pipeline because power estimations and the pattern transform induce an irreversible and non-linear transform at the beginning of this pipeline, and thus breaks the assumptions of MEG source modeling. Furthermore, single-trial power estimates are likely to be less contaminated in source space than in sensor space because beamforming can suppress noise (Litvak et al., 2010; Sekihara et al., 2004), and will thus help the linear decoder.

The performance of the source-first pipeline comes at a price, however: this approach induces a very large computational cost. For each trial, the MEG data must be projected onto a very large and redundant source space, and the power of each of its sources must then be estimated.

Our CSP-first pipeline alleviates this cost, performs comparably to the source-first pipeline, and reaches sharper source estimates. These results match our expectations: CSP identifies the few “virtual sources” (i.e., the few orthogonal components) that best discriminate the stimuli, implicitly estimates power, but remains linear. Critically, this approach proves to be ten times faster than the source-first pipeline.

We confirm these simulation results with real MEG data: the source-first and CSP-first pipeline perform comparably in decoding the perception of real versus scrambled faces using only high-frequency activity, and both outperform sensor space decoding. While the ground truth of these MEG signals is unknown, we can expect that the real and scrambled faces elicit different responses in visual cortex (Dobel et al., 2011; Zion-Golumbic and Bentin, 2007). The source estimates of the source-first pipeline and our CSP-first pipeline concur with this prediction: both peak around the same voxel locations in occipital cortex. Similarly to our simulation, the source estimates of the CSP-first pipeline appear sparser than those of the source-first pipeline. At this stage, however, it remains unclear whether this difference in activation spread reflects a true sparsity of the neural representations, or an uninformative bias of our method. Further work combining MEG and functional magnetic resonance imaging (fMRI) or intracranial recordings will be necessary to investigate this matter.

Taken together, our results unify (1) efficient models developed for sensor space decoding with (2) the interpretability of source space patterns. Our approach can thus help reduce the gap between the need to maximize signal-to-noise ratio, for e.g., brain-computer-interface objectives, and the necessity to remain interpretable, for e.g., neuroscientific theories.

## Supplementary material

**Figure S1:**
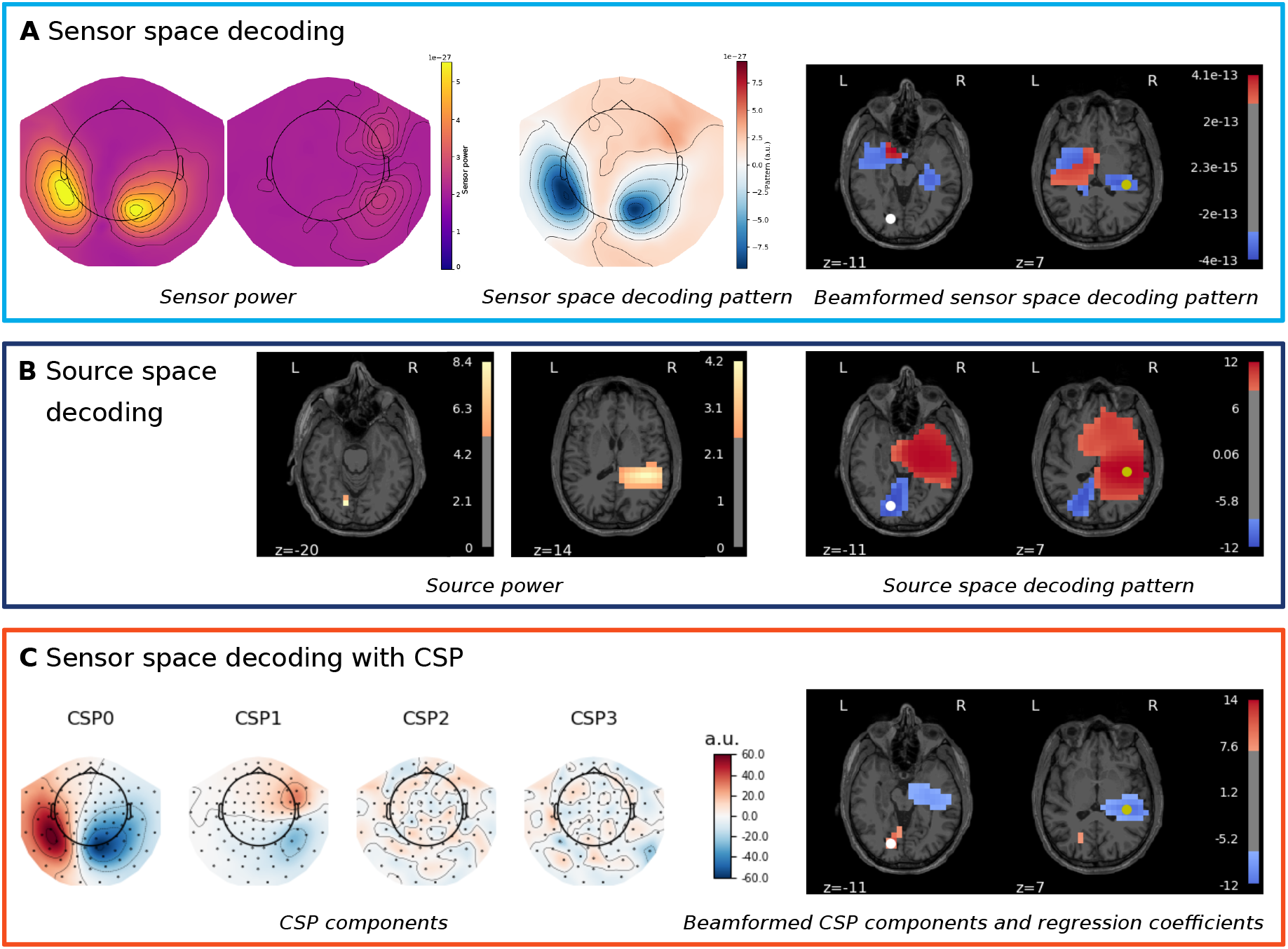
Visualization of decoding patterns for the simulation. The example corresponds to the *visual-audio* contrast and an input SNR of −10 dB. In all source plots, the true sources for the two classes are marked with a yellow and white dot. **A** Power-first pipeline. Sensor space power (left), sensor space decoding pattern after logistic regression (middle) and source space projection (right). **B** Source-first pipeline. Source space power (left) and source space decoding pattern (right). **C** CSP-first pipeline. CSP components (left) and the beamformed source estimate of the combined CSP components and logistic regression coefficients (right).

